# Low costs of adaptation to dietary restriction

**DOI:** 10.1101/2019.12.19.883058

**Authors:** Roy Z. Moger-Reischer, Elizabeth V. Snider, Kelsey L. McKenzie, Jay T. Lennon

## Abstract

Dietary restriction (DR) is the most successful and widespread means of extending organismal lifespan. However, the evolutionary basis of life extension under DR remains uncertain. The traditional evolutionary explanation is that when organisms experience DR, they allocate endogenous resources to survival and postpone reproduction until conditions improve. However, this life-extension strategy should be maladaptive if DR continues for multiple generations due to tradeoffs between longevity and reproduction. To test this prediction, we subjected the budding yeast *Saccharomyces cerevisiae* to 1,800 generations of evolution on restricted (i.e., DR) vs. non-restricted diets. Adaptation to a non-restricted diet improved reproductive fitness by 57% on that diet, but provided a much smaller (14%) advantage on a restricted diet. In contrast, adaptation to DR resulted in an approximately 35% increase in reproductive fitness on both restricted and non-restricted diets. Importantly, the life-extending effect of DR did not decrease following long-term evolution on the restricted diet. Thus, contrary to theoretical expectations, we found no evidence that the life-extending DR response became maladaptive during multigenerational DR. Our results suggest the DR response may have a low cost and that this phenomenon may have evolved for reasons that extend beyond the benefits of postponing reproduction.

## INTRODUCTION

Dietary restriction (DR) occurs when an organism experiences a reduction in energy consumption without any vitamin or mineral deficiencies [1]. It is the most consistent intervention known to extend lifespan, effective in taxa ranging from single-celled organisms to primates [1–6]. The conserved molecular mechanisms and genetic pathways by which DR promotes longevity have been well described [5,7]. Yet, the evolutionary basis of the life-extending DR response is not well understood [8] despite the fact that it has important implications for topics ranging from modern medicine to life-history theory [5,8–11].

The longstanding evolutionary explanation of the DR response is the postponed reproduction hypothesis, based on the disposable soma theory of aging [10,12–16]. Disposable soma theory describes a resource allocation tradeoff between somatic and reproductive functions. An organism can increase longevity by investing resources in the repair of somatic cell damage that accrues over time. Alternatively, esources can be allocated to the germline, which increases reproductive ability. Thus, lifespan could theoretically be extended by increasing allocation to the soma, for example in response to DR, although this should incur a tradeoff manifested as decreased reproduction.

Despite the tradeoff, life extension afforded by DR is thought to be adaptive in environments with fluctuating resource supply [10,12–15]. Specifically, the postponed reproduction hypothesis states that surviving periods of resource limitation via extended lifespan is beneficial because it allows individuals to reproduce later when conditions are more favorable [13–15]. If this hypothesis is correct, then organisms must outlive the duration of the poor conditions in order for extended lifespan and postponed reproduction to yield increased fitness. Otherwise, if resource limitation continues for multiple generations, there would be no payoff and the longevity-extending DR response should become maladaptive [10,17]. Therefore, multigenerational DR should favor individuals that cease to respond to DR with extended lifespan at the cost of allocation to reproduction.

In this study, we tested predictions about longevity and reproductive fitness in response to multigenerational DR by conducting long-term experimental evolution trials with the budding yeast, *Saccharomyces cerevisiae*. Budding yeast is a model organism in the study of aging, due to its easily measured lifespan and the fact that many mechanisms and genetic pathways involved in yeast aging are shared across eukaryotic taxa [5,7,8,18]. Although originally developed to describe aging in animals, the disposable soma theory applies readily to single-celled organisms such as *S. cerevisiae* that exhibit mother-offspring reproductive asymmetry (e.g., by budding) [19–21]. Allocation between longevity and reproduction, and adaptive responses to DR, are highly relevant in yeast life histories, because feast-and-famine cycles are typical for microorganisms in nature, including *S. cerevisiae* [22,23]. To test the postponed reproduction hypothesis, we selected for reproductive fitness for 1,800 generations. We measured life expectancy of evolved populations to assay for the expected diminishment of the DR response due to its predicted cost. We also measured evolved reproductive fitness to further investigate the effect of DR on adaptation.

## METHODS

### Evolution experiment

We used haploid (mating type *MAT*a) *Saccharomyces cerevisiae* of strain background W303, which is commonly used in aging research [24–26]. The experimental evolution was conducted using YPD-based non-restricted (NR: 10 g/L yeast extract, 20 g/L peptone, 20 g/L D-glucose) and restricted (DR: 5 g/L D-glucose) diets supplemented with 0.1 g/L adenine and 0.1 g/L tryptophan to address auxotrophies of the W303 strain. The 75% reduction in glucose concentration (20 g/L to 5 g/L) has been established to induce DR in *S. cerevisiae* [2,26]. We propagated five independent replicate lineages per treatment in 10 mL of medium by 1% v/v daily serial transfer for 275 days (>1,800 generations) in a shaking incubator (200 RPM) at 30 °C. Because there is little or no aging-associated mortality until the third day [18,26], this regime selects for reproductive fitness and does not select for longevity.

### Measurement of longevity

We measured longevity using a chronological lifespan (CLS) assay. This assay involves plating aliquots of post-stationary phase yeast cultures to measure survivorship over time. We grew yeast cultures in SDC medium (20 g/L or 5 g/L D-glucose, 6.7 g/L yeast nitrogen base, 0.08 g/L *myo*-inositol, 0.08 g/L uracil, 0.04 g/L PABA, 0.18 g/L adenine, 0.18 g/L tryptophan, 0.16 g/L leucine, 0.08 g/L all other amino acids) which greatly reduced the variation in lifespan measurements compared to YPD medium [18]. The concentrations of glucose (20 g/L or 5 g/L) corresponded with non-restricted (NR) and restricted (DR) conditions, respectively. Replicate aliquots from each cryopreserved evolved population were grown to stationary phase in NR or DR SDC medium for three days. Mortality was assumed negligible during this period [18]. To quantify the abundance of viable individuals, we counted colony-forming units after spread-plating dilutions onto non-restricted YPD agar plates and incubating them for three days at 30 °C. We sampled each culture in this fashion every two days until survivorship fell below 1% (NR: 16 days; DR: 38 days).

We quantified longevity by calculating the life expectancy (*e*_0_) from the life table derived from the CLS assay [27]: 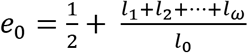, where *l*_0_, *l*_1_,… *l_ω_*, are the proportions of individuals surviving to timepoints 0, 1,…, *t*_final_[27]. We chose this metric because it is effective for comparing expected longevity between populations with the same life history and magnitude of lifespan [28].

### Measurement of reproductive fitness

We measured reproductive fitness by competing each evolved population against a common third-party strain in a competitive growth assay that integrated reproductive fitness over a 24-hr period [29]. We used strain YDL185W from the green fluorescent protein (GFP) clone collection (Invitrogen), which expresses Vma1p-GFP fusion protein [30], as the third-party yeast strain. We used the 488 nm laser of a Novocyte flow cytometer (ACEA Biosciences) to differentiate between GFP-expressing and non-GFP cells, and monitored changes in abundance of the two cell types over 24 hr. Relative reproductive fitness (*W*) versus the third-party was calculated as

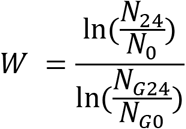

where *N*_0_ represents initial abundance of the focal strain, *N*_24_ is the abundance of the focal strain after 24 hr, and *N*_*G*24_ and *N*_*G*0_ are final and initial abundances of the Vma1p-GFP third-party strain [29]. We then expressed the reproductive performance of evolved lines on each diet relative to the W303 ancestor. To do this, we relativized *W* to the reproductive fitness of ancestor strain: 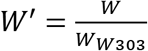, where *W*_W303_ is reproductive fitness of the W303 ancestor, calculated as before.

### Statistical analyses

We used two-way ANOVA to test for main effects of the evolution diets (NR vs. DR) and assay diets (NR vs. DR) on longevity and reproductive fitness, as well as their interaction. This model allowed us to evaluate yeast performance under both “home” (e.g., DR-evolved lines on DR diet) and “away” (e.g., DR-evolved lines on NR diet) conditions. We used post-hoc Tukey’s HSD tests to obtain adjusted *P*-values for pairwise comparisons. We compared the life expectancy of evolved lines to the ancestor using one-sample *t*-tests. All statistical analyses were conducted in *R* [31].

## RESULTS

### Longevity

As expected, lifespan was longer when yeast were cultured under dietary restriction (DR), confirming the DR phenomenon was successfully induced in our experiments. Prior to evolution, life expectancy of ancestral W303 was 75% higher on DR than non-restricted (NR) diet (8.4 ± 0.8 days and 14.7 ± 2.5 days, respectively; *t*_13_ = 2.80, *P* = 0.008). Among evolved lines, life expectancy was also extended on DR relative to NR, suggesting the DR response was retained during evolution (two-way ANOVA *F*_1,16_ = 21.4, *P* = 2.8 × 10^−4^) (figure 1). In contrast to predictions, life expectancy of DR-evolved lines was not shorter than the ancestor on DR (one-sample *t*_4_ = 0.36, *P* = 0.632). On an NR diet, life expectancy among evolved lines was marginally decreased relative to the ancestor (DR-evolved: one-sample *t*_4_ = −2.73, *P* = 0.053; NR-evolved: one-sample *t*_4_ = −2.30, *P* = 0.083). Finally, and again contrary to predictions, there was no significant difference in life expectancy between DR-evolved and NR-evolved lines (two-way ANOVA *F*_1,16_ = 0.699, *P* = 0.415).

**Figure 1.**
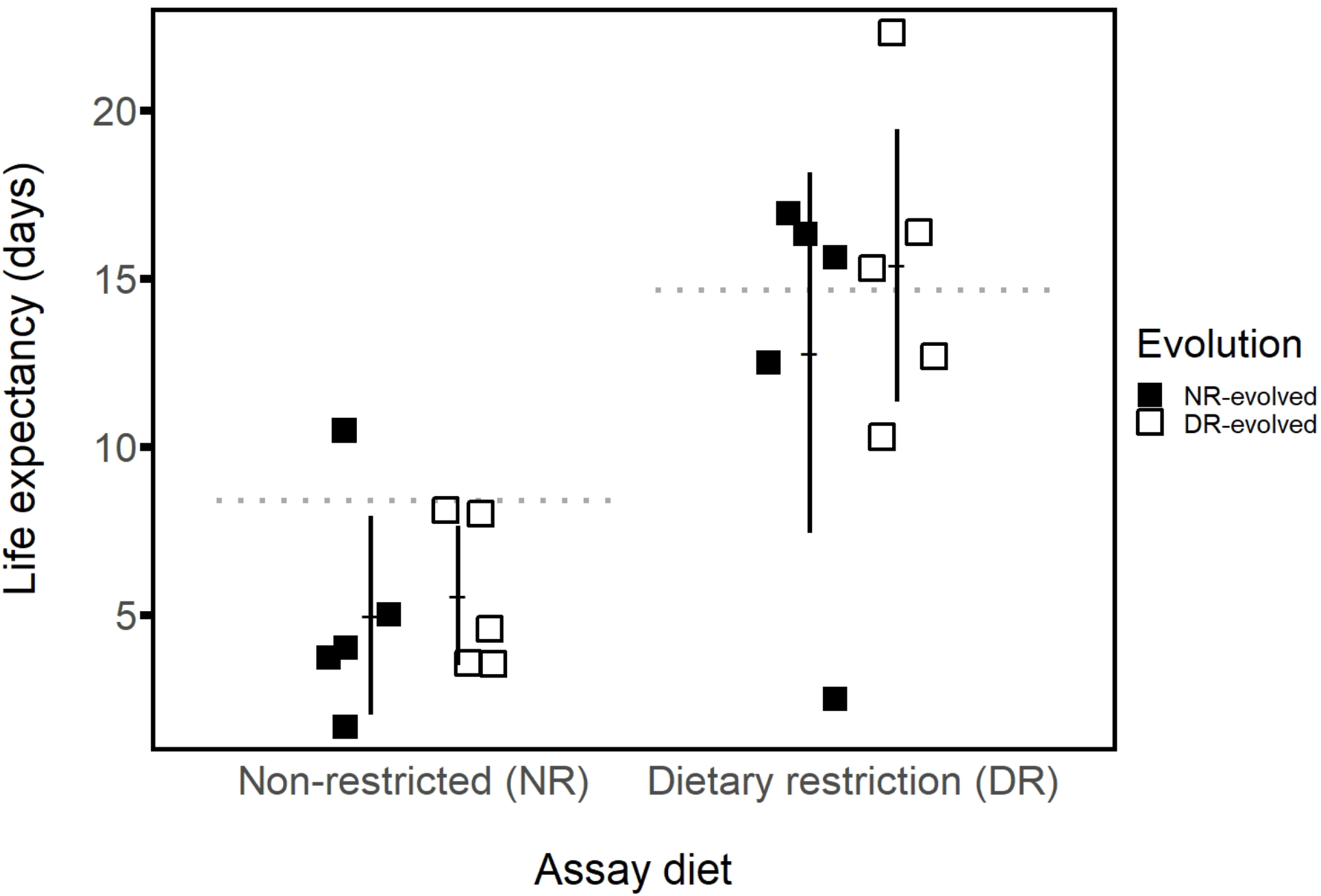
Life expectancy of non-restricted (NR)-evolved and dietary restriction (DR)-evolved budding yeast populations. The life-extending DR response was observed in evolved lines, which exhibited significantly longer life expectancy on DR. There were no decreases in life expectancy on DR relative to the ancestor (right dashed line). There were marginally significant decreases in life expectancy on NR relative to the ancestor (left dashed line). Life expectancy did not differ between the evolution treatments. Error bars: ± 2 standard errors of the mean.

### Reproduction

Evolved lines exhibited higher reproductive fitness on their “home” evolutionary diets than on their “away” evolutionary diets (two-way ANOVA *F*_1,16_ = 13.74, *P* = 0.002) (figure 2). However, pairwise comparisons revealed that the magnitude of this effect was not equal on either home diets or away diets. Specifically, lines evolved on NR exhibited a 19% higher reproductive fitness on their home diet than did DR-evolved lines on their home diet (*P_adj_* = 0.007). When assayed on the away diet, reproductive fitness was 24% higher for the DR-evolved lines assayed on NR diet than for NR-evolved lines assayed on DR (*P_adj_* = 0.003). The reproductive fitness of the DR lines was the same whether they were assayed on the home or away diet (*P_adj_* = 0.488), while the reproductive fitness of the NR-evolved lines was 27% lower on their away diet (*P_adj_* = 2.80×10^−5^).

**Figure 2.**
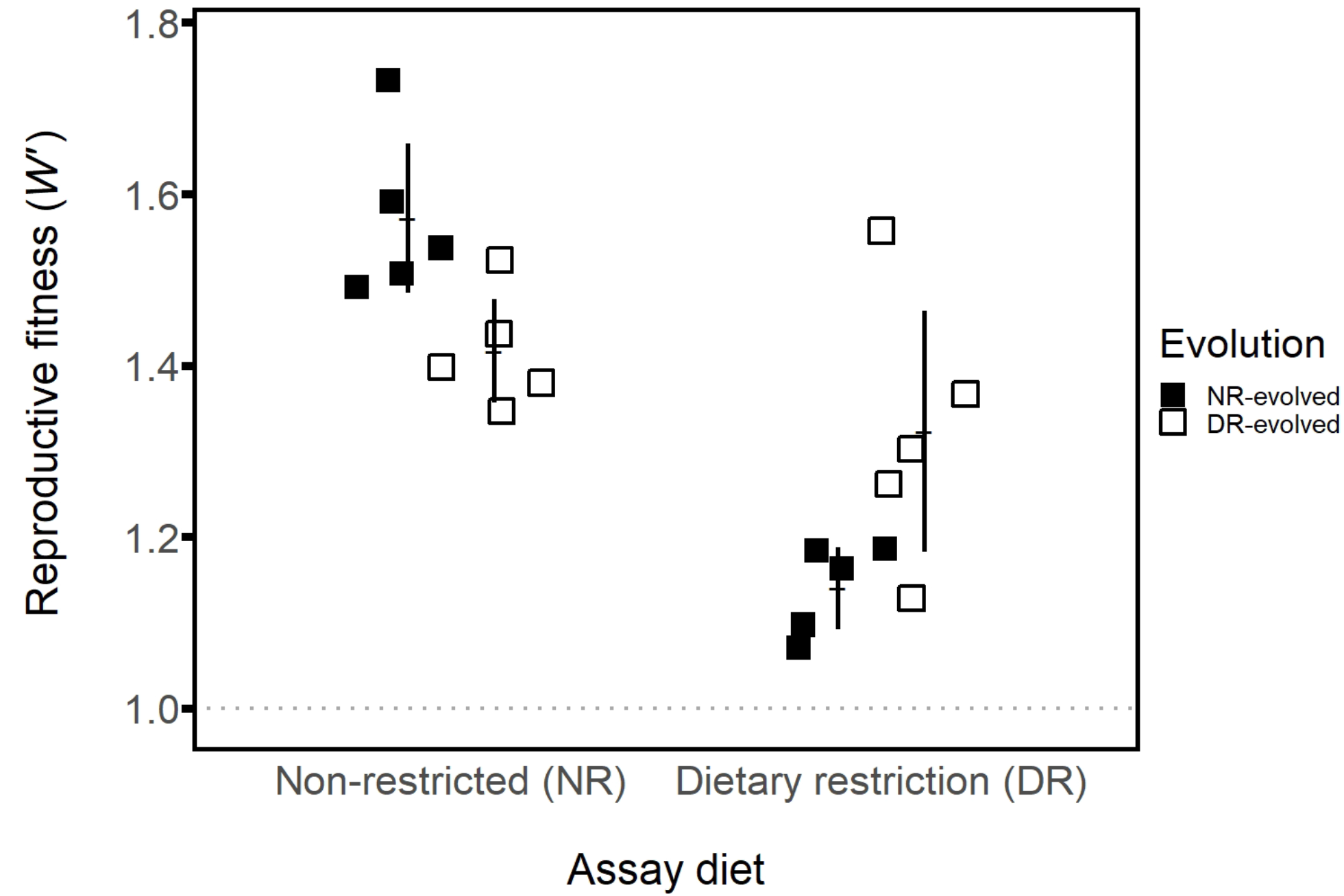
Reproductive fitness of evolved budding yeast lines relative to ancestral reproductive fitness (dashed line). Reproductive fitness increased across all environments and was higher on “home” evolutionary diets than on “away” diets. Non-restricted (NR)-evolved lines attained higher reproductive fitness on their home diet than dietary restriction (DR)-evolved lines did on theirs. Meanwhile, lines evolved on DR performed better on their away diet than did NR-evolved lines. In fact, the reproductive fitness of DR-evolved lines was not significantly different on the away diet versus the home diet. Meanwhile, NR-evolved lines exhibited poor performance on their away diet. Errors bars: ± 2 standard errors of the mean.

## DISCUSSION

Our results from a long-term evolution experiment with budding yeast do not support the postponed reproduction hypothesis for the evolution of the life-extending response to DR. Specifically, the DR response was not diminished in yeast populations that evolved under long-term DR and there was no difference in life expectancy between the DR-evolved and NR-evolved populations regardless of assay condition (figure 1). Together, these findings suggest the DR response was not maladaptive when the organisms were challenged by resource limitation for multiple generations. Instead, the evolutionary retention of the DR response suggests it may be adaptive irrespective of a payoff from postponed reproduction.

We also found that home-diet reproductive fitness of the NR-evolved populations was higher than that of DR-evolved populations. This finding too is contrary to the postponed reproduction hypothesis. According to the postponed reproduction hypothesis, the DR response is the optimum phenotype [32] in fluctuating resource environments [10,13,14]. In a constant environment, however, the optimum phenotype is one with limited longevity in order to attain high reproduction. This means that in our laboratory evolution with constant diet, and no selection for longevity, the long-lived DR phenotype would be farther away from the optimum than the shorter-lived non-restricted phenotype. Because of the increased distance to the phenotypic optimum, reproductive fitness is predicted to increase by a greater amount during evolution on DR [32]. However, we observed the opposite, suggesting the DR response phenotype was not farther from the optimum.

Evidence from other systems has also cast doubt on the universality of the postponed reproduction hypothesis. For example, in experimental evolution trials with *Drosophila melanogaster*, longevity was comparable for populations irrespective of the diets on which they evolved [17]. In a separate study, diet fluctuations within the lifespans of *D. melanogaster* individuals demonstrated no advantage to postponed reproduction [34]. Specifically, there was high mortality and no increase in fecundity when flies were switched from restricted to non-restricted diet, suggesting the absence of postponed payoff [34]. Thus, our study is consistent with observations made in a separate model system and suggests that other mechanisms can contribute to the evolution of extended life span under DR.

One potential explanation for our findings is that evolution on DR confers other advantages. For example, the results suggest that maintaining populations on a restricted diet led to the evolution of better generalists. Specifically, DR-evolved lines had higher reproductive fitness on the away diet than did NR-evolved lines on the away diet. Moreover, reproductive fitness of DR-evolved lines was not significantly lower on the away diet than the home diet (figure 2). Thus, adaptation to dietary stress resulted in the evolution of generalists. Interestingly, this bears similarity to other findings which have shown that generalist microbial species tend to be better adapted to a different type of stress, namely desiccation [33]. Our study’s DR-evolved generalists also align with experimental evolution trials that used *D. melanogaster*, where DR-evolved lines exhibited high reproductive fitness on all assay diets tested [17].

Together, our results show that the DR response did not become maladaptive after multigenerational DR. This suggests that the DR response may be adaptive by means other than postponed reproduction, perhaps having evolved as part of a more general environment-responsive adaptation than a response to nutrition status alone [8]. Second, we observed low costs of adaptation to DR. DR-evolved lines exhibited improved reproductive fitness on DR, a better generalist strategy than NR-evolved lines, and no tradeoff in the ability to extend lifespan via DR. The low cost of DR specialization in our study suggests that phenotypic plasticity in response to nutrition status may not have evolved solely due to constraints of resource allocation [8].

## ACKNOWLEDGEMENTS

We thank Soni Lacefield for providing ancestor yeast strains and Farrah Bashey-Visser for critical feedback and suggestions.

